# A novel, fig-associated microbe promotes reproductive success via variable life history mechanisms in *C. elegans* and *C. inopinata*

**DOI:** 10.64898/2025.12.16.694684

**Authors:** Austin C. Link, Kimberly A. Moser, John Wang, Gavin C. Woodruff

## Abstract

Variation in life history strategies is among the most striking features of animal diversity. Simultaneously, the microbes an animal interacts with are a critical and dynamic aspect of the host environment that can have profound impacts on their life history traits. As microbial environments diverge across animal lineages, life histories and their responses to such microbial contexts are expected to evolve as a consequence. *Caenorhabditis* nematodes are bacterivores that exhibit a diversity of life history strategies and fill diverse ecological niches. *C. elegans* thrives on rotting plants and grows rapidly with high fecundity; *C. inopinata* thrives in fresh figs and grows more slowly with lower fecundity. To understand how hosts with divergent life histories and ecologies respond to the microbes they interact with, we isolated over forty bacterial species from the natural fig environment of *C. inopinata*. This microbial survey revealed an isolate, *Klebsiella* sp. WOUb2, that doubles the population growth rate of *C. inopinata*. While this isolate also increases the population growth rate of *C. elegans, Klebsiella* sp. WOUb2 increases individual fecundity and developmental rate in *C. elegans*, whereas it only increases developmental rate of *C. inopinata*. Thus, fitness is modulated by variable life history mechanisms in the two species. Comparisons with nucleotide databases reveal *Klebsiella* sp. WOUb2 is closely related to other *Klebsiella* isolates known to influence *Caenorhabditis* nematode fitness. Additionally, the similarity of *Klebsiella* sp. WOUb2 to microbes associated with fig wasps and figs suggests *C. inopinata* frequently encounters this microbe in its natural context. Taken together, this shows that different physiological responses can underlie conserved, beneficial interspecific interactions.

## Introduction

Life history strategies vary tremendously across species. For instance, an oak tree produces hundreds of thousands of seeds over a lifetime of hundreds of years (Greenberg 2000; Guyette et al. 2004; Drobyshev et al. 2008; Caignard et al. 2017). Conversely, albatrosses rear less than one fledged chick, on average, per year across a lifetime of decades (Weimerskirch et al. 1987). And fruit flies produce hundreds of eggs in a lifetime of weeks (Flatt 2020). Clearly, vastly different reproductive ways of life have evolved. Indeed, evolutionary theory predicts the direction of life history evolution is contingent on the current population state (Stearns 1992). That is, in some cases, faster development maximizes fitness whereas in others, extending the reproductive duration is more effective (Stearns 1992). Not only is historical contingency predicted to impact theoretical life history outcomes— long-term environmental changes often influence fundamental life history traits such as fecundity, survival, and developmental rate (Coulson et al. 2011; Lancaster et al. 2017). Indeed, understanding how evolutionary history and environmental change interact to produce the enormous diversity in life history strategies across the tree of life is a fundamental goal of evolutionary biology.

One dynamic component of an organism’s biotic environment is the myriad number of microorganisms it interacts with. Microbes are ubiquitous and frequently influence organismal phenotypes, including life history traits (Engelstädter and Hurst 2009; Than et al. 2020; Henry et al. 2021). For instance, bacterial pathogens can decrease fecundity in numerous animals including *Daphnia* crustaceans (Vale and Little 2012) and *Drosophila melanogaster* flies (Kutzer and Armitage 2016); different microbiomes can modulate flowering time in *Arabidopsis* (Panke-Buisse et al. 2015); and native microbiomes are critical for woodrat survival (Kohl et al. 2014). Moreover, as environments change, microbial communities are expected to change and evolve as a consequence (Hutchins et al. 2019; Jansson and Hofmockel 2020; Trivedi et al. 2022). Thus, there is ample opportunity for species to diverge in differing coevolving microbial environments. How do evolving host/microbe interactions influence life history traits?

Understanding the coevolution of host/microbe interactions requires the appropriate model systems. The nematode *Caenorhabditis elegans* is a long-standing model for molecular and developmental genetics (Corsi et al. 2015). Associated with rotting plant detritus (Frézal and Félix 2015; Schulenburg and Félix 2017), *C. elegans* is frequently found with and consumes a diverse array of bacterial species in its natural environment (Zhang et al. 2017). Moreover, multiple bacterial strains have been isolated from the natural environment of *C. elegans*, and these strains promote both beneficial and deleterious life history responses (Dirksen et al. 2016; Samuel et al. 2016; Zhang et al. 2021). Together with the isolation of *Caenorhabditis* species with their own associated microbial communities, this group offers an ideal system for disentangling host/microbe interactions and their influence on the evolution of life history strategies.

*C. inopinata* is a sister species of *C. elegans* (Kanzaki et al. 2018) with a divergent natural environment (Kanzaki et al. 2018; Woodruff and Phillips 2018) and life history strategy (Kanzaki et al. 2018; Woodruff et al. 2019). Whereas *C. elegans* thrives on rotting plants, *C. inopinata* reproduces in fresh figs (Kanzaki et al. 2018; Woodruff and Phillips 2018). And whereas *C. elegans* harbors rapid growth and high fecundity (Byerly et al. 1976), *C. inopinata* exhibits slow development (double the rate of *C. elegans*) and low fecundity (an order of magnitude lower than *C. elegans*) (Woodruff et al. 2019). These species also differ in reproductive mode— *C. elegans* populations harbor mostly self-fertile hermaphrodites with rare males (Cutter et al. 2019); *C. inopinata* is a female/male species (Kanzaki et al. 2018). Because of their divergent ecological environments and life histories, the *C. elegans*/*C. inopinata* sister species pair represent a solid contrast for understanding the relationship between divergence, life history traits, and host/microbe interactions. Indeed, previous studies have revealed hints of a conserved relationship with *Enterobacteriaceae* bacteria across *Caenorhabditis* species. For example, *Raoultella* sp. JUb54, isolated from rotting apples, promotes increased population growth in *C. elegans* (Samuel et al. 2016; Bubrig et al. 2020), and *Raoultella* sp. CiN001 promotes increased population growth in *C. inopinata* (Kanzaki et al. 2018). Here, we probe the life history mechanisms underlying this microbe-driven increase in reproduction. Specifically, we find that a likely ancient relationship with *Enterobacteriaceae* bacteria promotes fitness via variable life history mechanisms in *C. elegans* and *C. inopinata*.

## Results

### A survey of fig microbes reveals bacteria that modulate C. inopinata fitness

To discover the natural microbes living with *C. inopinata*, we first collected 38 fresh figs from 12 *F. septica* plants (Woodruff et al. 2024). We then isolated live bacterial monocultures from the inside of the figs, successfully establishing 45 isolates (Fig. 1A-B). Sanger sequences of the 16S locus revealed the vast majority of these belonged to the Gammaproteobacteria class (Fig. 1B; 33/45 isolates), with the rest belonging to other groups (from the Actinobacteria, Actinomycetota, Bacteroides, and Firmicutes phyla; Supplemental Table Sheet 1).

**Figure 1.**
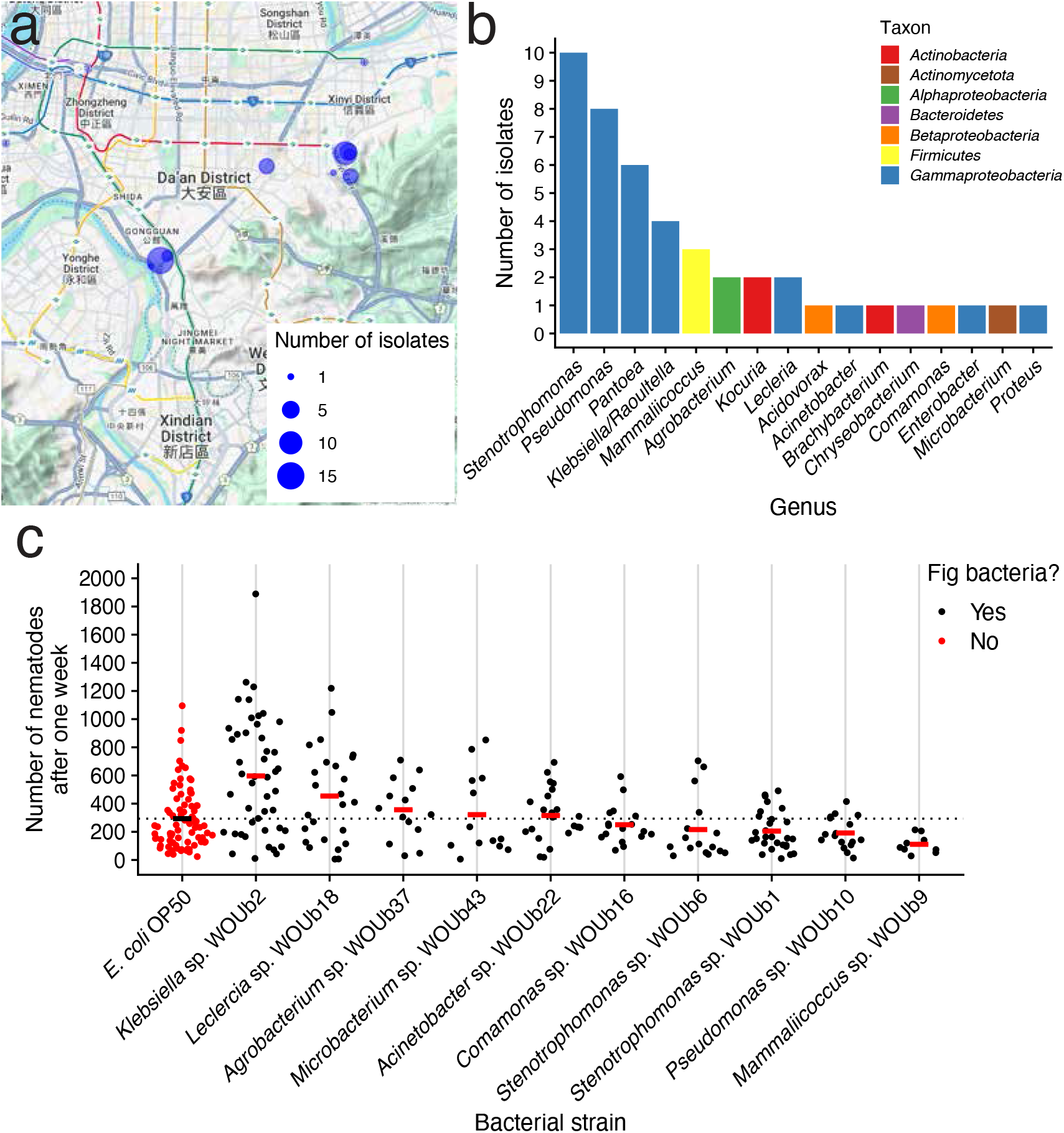
*C. inopinata* responds to bacteria isolated from fresh *Ficus septica* figs. (a) Microbes were isolated from figs in Taipei, Taiwan. The sites of individual *F. septica* plants where figs were sampled are shown as blue points which are sized by the number of microbial strains isolated from each plant. Maps were generated in part with Google Maps and their data sources. © 2024 Google (https://about.google/brand-resource-center/products-and-services/geo-guidelines/). (b) The number of microbial strains per genera among our 45 strains isolated from figs. (c) *C. inopinata* shows variable population growth when reared on bacteria from the fig environment. Each data point represents the number of nematodes on a single plate (founded by one female and four males) at the end of the seven-day period for a given bacterial strain. Crossbars represent means; the horizontal dotted line represents the *E. coli* OP50 mean. Sina plots are jittered scatterplots that take the contours of a violin plot. N_crosses_ = 10-80.

To determine the impact of the natural microbes on nematode fitness, we assayed *C. inopinata* growth when reared on a subset of these strains and *E. coli* OP50 (the laboratory standard *C. elegans* bacterial food (Stiernagle 2006)). To assay population growth, we placed one virgin female with four males and counted the total number of nematodes on that plate a week later, which corresponded to 2-3 generations of nematode proliferation. Indeed, *C. inopinata* population growth varies by microbial food source (Fig. 1c). However, only one fig-associated microbe increased nematode population growth compared to *E. coli* OP50— *Klebsiella* sp. WOUb2 (Fig. 1c; 115% increase; post-hoc Tukey test adjusted *p* < 0.001; Cohen’s *d* = 0.82). Thus, at least one microbe originating from the natural environment of *C. inopinata* is capable of dramatically improving nematode population growth rates compared to conventional laboratory conditions, consistent with previous studies in *C. elegans* (Dirksen et al. 2016; Samuel et al. 2016).

*A microbe that promotes divergent life history responses in C. elegans and C. inopinata* Population growth rates emerge from the interaction of several life history traits such as fecundity, viability, and developmental rate (Stearns 1992). To understand how *Klebsiella* sp. WOUb2 drives increased population growth rate in nematodes, we measured these life history traits in *C. inopinata* (Fig. 2, Supplementary figures 2-4). In addition, we looked at the life history responses of *C. elegans* to this microbe to understand how bacteria-associated life history responses might evolve among divergent species. As *C. inopinata* and *C. elegans* differ in reproductive mode, we focused on *C. elegans fog-2(q71)* mutants for these comparisons (while also reporting observations for wild-type *C. elegans* PD1074 animals). Homozygous *fog-2* mutants inhibit hermaphrodite spermatogenesis and transform *C. elegans* into an obligate outcrosser (Schedl and Kimble 1988)). Finally, to test for potential temperature effects, we also conducted these experiments at two temperatures, 20°C and 25°C.

**Figure 2.**
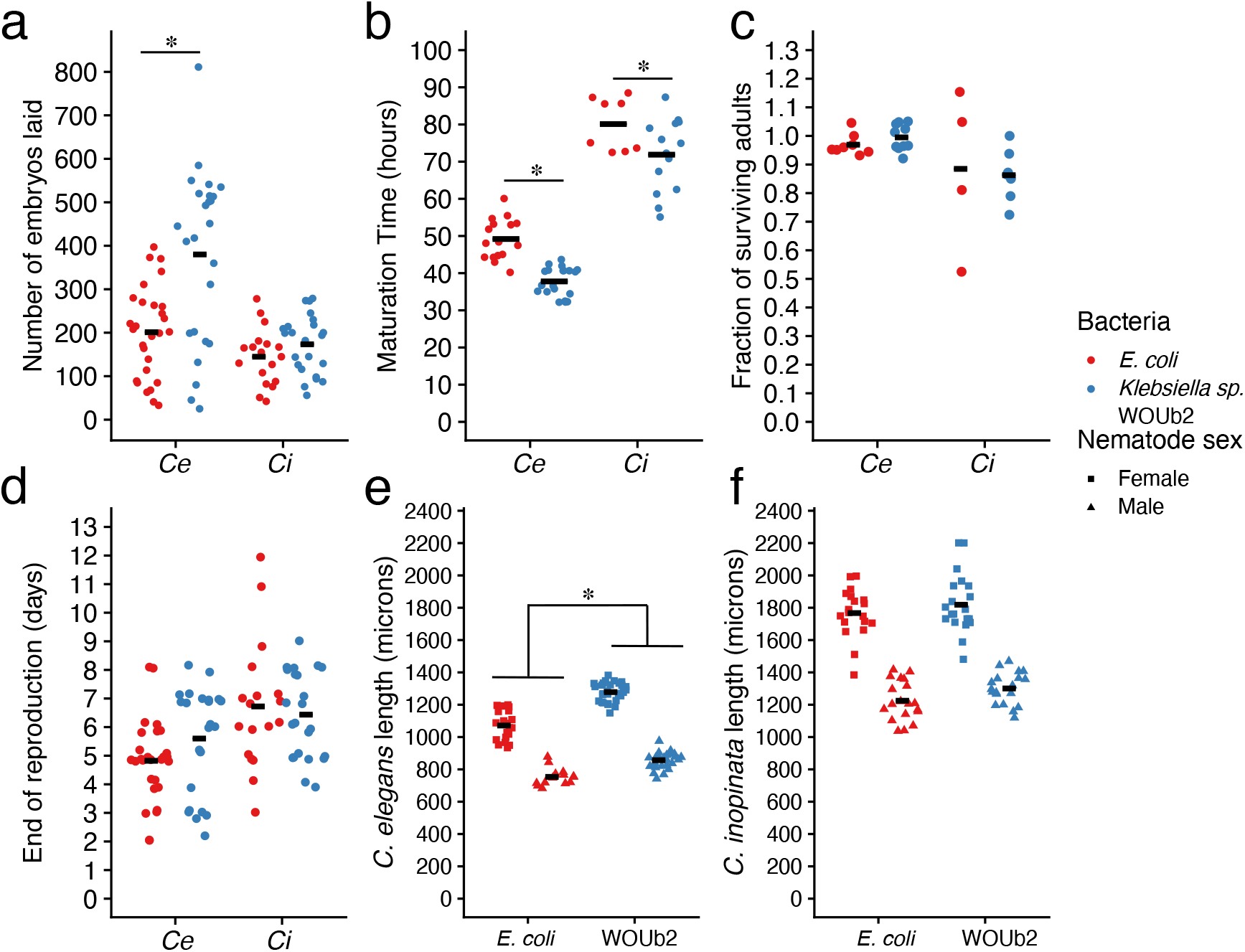
*Klebsiella* sp. WOUb2 promotes divergent life history responses in *C. inopinata* and *C. elegans*. (a) Individual fecundity. *Klebsiella* sp. WOUb2 promotes higher individual fecundity in *C. elegans*, whereas in *C. inopinata*, individual fecundity is not changed. Each point represents the total number of embryos produced by an individual female. (b) Developmental timing. *Klebsiella* sp. WOUb2 promotes faster development in both *C. inopinata* and *C. elegans*. Each point represents the median time to adulthood for a given plate. (c) Viability. Embryo-to-adult viability is unaffected for both *C. elegans* and *C. inopinata* in all rearing conditions. Each point represents the fraction of embryos surviving to adulthood. Fractions above 100% are likely due to undercounting the starting number of embryos. (d) Reproductive period. *Klebsiella* sp. WOUb2 does not promote any major differences in reproductive duration among species. Each point represents the age of the last day of reproduction of an individual female. *(e) C. elegans* body size. *Klebsiella* sp. WOUb2 promotes increased body size in *C. elegans* adults. Each point represents the length of a one-day old adult. *(f) C. inopinata* body size. *Klebsiella* sp. WOUb2 and *E. coli* promote comparable body sizes in *C. inopinata* adults. Each point represents the length of a one-day old adult.

*Klebsiella* sp. WOUb2 also increased the population growth rate of *C. elegans* (Supplemental Figure 1; Wild-type *C. elegans*: 41% increase, Cohen’s *d* = 0.78, Wilcoxon rank-sum test *W*=384, *p*=0.0076, N=29-30; *C. elegans fog-2*: 95% increase, Cohen’s *d* =1.4, Wilcoxon rank-sum test *W*=441, *p*=0.00028, N=18). As *Klebsiella* sp. WOUb2 increased the population growth rate of both species, we likewise hypothesized this species would increase individual fecundity in both nematode species. Surprisingly, *Klebsiella* sp. WOUb2 had no detectable impact on the lifetime reproductive output of individual *C. inopinata* females irrespective of rearing temperature (Fig. 2a and Supplemental Figure 2a; post-hoc Tukey test adjusted *p* > 0.05). Conversely, *Klebsiella* sp. WOUb2 increased individual fecundity compared to *E. coli* in *C. elegans fog-2(q71)* mutants at both 20°C (Supplemental Figure 2a; Cohen’s *d* = 1.882, post-hoc Tukey test adjusted *p* < 0.001) and 25°C (Fig. 2a; Cohen’s *d* = 1.141, post-hoc Tukey test adjusted *p* < 0.001). For self-fertile *C. elegans* hermaphrodites, there was no difference in individual reproductive output between those grown on *Klebsiella* sp. WOUb2 and *E. coli*, irrespective of temperature (Supplemental Figure 3a-b; post-hoc Tukey test adjusted *p* > 0.05). Consequently, *Klebsiella* sp. WOUb2 must promote *C. inopinata* population growth (Fig. 1c) via mechanisms not related to individual reproductive output.

**Figure 3.**
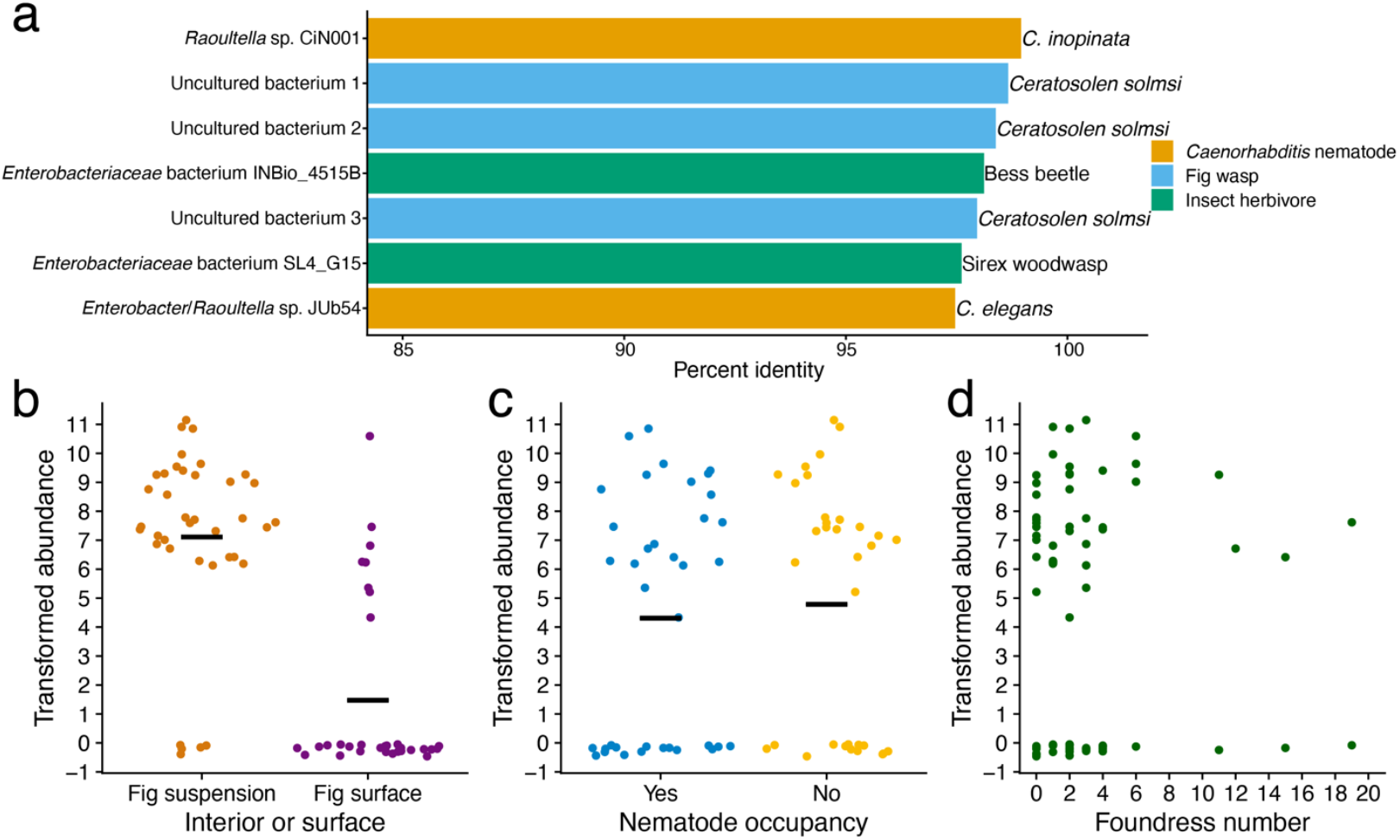
*Klebsiella* sp. WOUb2 is frequently encountered by *C. inopinata* in the wild. (a) The 16S sequence of *Klebsiella* sp. WOUb2 is similar to other microbes associated with *Caenorhabditis* nematodes, fig wasps, and other insect invertebrates. Reported are the top seven BLAST hits to the NCBI nr database by percent identity. *Raoultella* sp. CiN001 was previously isolated from *C. inopinata* (Kanzaki et al. 2018). Three uncultured bacteria associated with *Ceratosolen solmsi* pollinating fig wasps are included (accession numbers HQ639492, HQ639491, and HQ639545). Two insect-associated hits are included (*Passalidae* bess beetle (Vargas-Asensio et al. 2014); *Sirex noctilio* woodwasp (Adams et al. 2011)). *Enterobacter* sp. JUb54 was originally isolated from rotting apples harboring *C. elegans* (Samuel et al. 2016) and is likely a member of the genus *Raoultella*/*Klebsiella* (Bubrig et al. 2020). (b-d) An ASV with 100% identity to *Klebsiella* sp. WOUb2 was identified in multiple *Ficus septica* fig samples connected to a 16S metabarcoding study (Woodruff et al. 2024). Here, center log ratio-transformed abundances of this specific ASV are plotted. For (b-c), crossbars represent means. Sina plots are strip plots that take the contours of a violin plot. (b) The *Klebsiella* sp. WOUb2 ASV is more abundant in fig interiors than on fig surfaces. (c) The abundance of the *Klebsiella* sp. WOUb2 ASV is not impacted by the presence of nematodes. (d) The abundance of the *Klebsiella* sp. WOUb2 ASV does not covary with the number of pollinating fig wasp foundresses.

Populations that mature faster grow faster (Stearns 1992), so next, we examined the impact of *Klebsiella* sp. WOUb2 on nematode developmental rates (Fig. 2b; Supplemental figure 2b). In For all plots, crossbars represent means. Sina plots are jittered scatterplots that take the contours of a violin plot.contrast to its impact on fecundity, *Klebsiella* sp. WOUb2 caused *C. inopinata* animals to mature 8 hours faster in comparison to *E. coli* when reared at 25°C (Fig. 2b; Cohen’s *d* = 0.914, post-hoc Tukey test adjusted *p* < 0.05) but not at 20°C (Supplemental Figure 2b). Additionally, *Klebsiella* sp. WOUb2 had a detectable impact on the *C. elegans fog-2(q71)* mutants at 25°C (*fog-2(q71)*: mean difference of 11 hours, Cohen’s *d* = 2.452, post-hoc Tukey test adjusted *p* < 0.001).

Fecundity and developmental rate are not the only life history traits that contribute to reproductive success (Stearns 1992). To better describe nematode life histories, measures of viability, reproductive duration, and body size were taken. We found that neither *Klebsiella* sp. WOUb2 nor *E. coli* affected embryo-to-adult viability in either nematode species (Fig. 2c, Supplemental Figure 2c; post-hoc Tukey test adjusted *p* > 0.05). These bacteria likewise had no effect on reproductive duration (Fig. 2d; Supplemental Figure 2d). *Klebsiella* sp. WOUb2 promoted increased adult body sizes in *C. elegans* (Fig. 2e; 14% increase (*fog-2(q71)* males), 19% increase (*fog-2(q71)* females); Cohen’s *d* = 2.0 (males), 2.9 (females); Wilcoxon rank-sum test *W*=328 (males), 600 (females); both *p*<0.0001, N=14-29), while having no obvious impact on body size in *C. inopinata* (Fig. 2f; 6% increase (males), 3% increase (females); Cohen’s *d* = 0.69 (males), 0.31 (females); Wilcoxon rank-sum test *W*=240 (males), 218 (females); *p*=0.085 (males), 0.63 (females); N=19-21). Taken together, *Klebsiella* sp. WOUb2 promotes variable life history responses in *C. elegans* and *C. inopinata*, despite increasing population growth rates in both species.

*Klebsiella species are frequently encountered by C. inopinata in its natural context* Because of its diverse impact on nematode life history responses (Fig. 2), we sought to better understand the evolutionary and ecological context of *Klebsiella* sp. WOUb2. We assembled the genome of *Klebsiella* sp. WOUb2 (Miller et al. manuscript in prep) and extracted the complete 16S rRNA sequence to identify similar bacterial species in the NCBI nr database (Camacho et al. 2009). The top BLAST hits revealed *Klebsiella* sp. WOUb2 has high sequence identity to bacteria previously associated with *Caenorhabditis* nematodes, fig wasps, and plant-associated insects (Fig. 3a). The sequence with the highest percent identity was that of a microbe previously isolated from *C. inopinata* (*Raoultella* sp. CiN001; 99.0% identity; Supplemental Table Sheet 2) (Kanzaki et al. 2018). Notably, this isolate also increased the *C. inopinata* population growth rate (Kanzaki et al. 2018). Additionally, the seventh-highest hit was *Raoultella* sp. JUb54 (Fig. 3a), isolated from rotting apples associated with *C. elegans* (Samuel et al. 2016; Bubrig et al. 2020). This strain is beneficial to *C. elegans* and can increase its population growth rate (Samuel et al. 2016). In addition, three top hits included those associated with an unpublished study of the microbiome of the pollinating fig wasp, *Ceratosolen solmsi* (Fig. 3a). *C. solmsi* is in the same genus as the pollinator of *F. septica* (*C. bisulcatus* (Yokoyama and Iwatsuki 1998; Xiao et al. 2013)), and *Caenorhabditis* nematodes have also been found in association with *C. solmsi* (Jauharlina et al. 2022). The fourth- and sixth-highest BLAST hits were sequences associated with *Passalidae* bess beetles (Vargas-Asensio et al. 2014) and *Sirex noctilio* woodwasp larvae (Adams et al. 2011), respectively (Fig. 3a); both are plant-associated herbivores. Most of the other BLAST hits were associated with less similar, human-associated *Klebsiella* species (Supplemental Table Sheet 2). This suggests *Klebsiella* sp. WOUb2 is closely related to bacteria frequently found in environments similar to those encountered by *C. inopinata* (e.g., *C. inopinata* and fig wasp sample BLAST hits) and *Caenorhabditis* nematodes more broadly (e.g., *C. elegans* and plant-associated insect BLAST hits; Fig. 3a).

We previously described the microbial communities of *F. septica* figs (Woodruff et al. 2024). To further explore the connection between *Klebsiella* sp. WOUb2 to the natural environment of *C. inopinata*, we aligned the 16S sequences from that metabarcoding project to the *Klebsiella* sp. WOUb2 genome. Subsequently, we discovered an Amplicon Sequence Variant (ASV) with 100% sequence identity to *Klebsiella* sp. WOUb2. This ASV was highly abundant in our fig samples (Fig. 3b-d). Because we collected additional field data with these samples (Fig. 3b-d) (Woodruff et al. 2024), were able to make additional inferences about the ecology of this ASV. The *Klebsiella* sp. WOUb2 ASV was more abundant in fig suspensions compared to fig surface washes (Fig. 3b; Wilcoxon rank-sum test *p* < 0.001, W = 1074), suggesting this species is more commonly found in fig interiors. Conversely, we found no association between the abundance of the *Klebsiella* sp. WOUb2 ASV and nematode occupancy (Fig. 3c; Wilcoxon rank-sum test *p* = 0.51, W = 554) nor with the number of pollinating fig wasp foundresses in a given fig (Fig. 3d; OLS linear model *p* = 0.85, *F* = 0.036). Thus, while *Klebsiella* sp. WOUb2 is likely frequently found in the lumen of *F. septica* figs (Fig. 3b), its abundance does not appear to depend on nematode presence (Fig. 3c) nor pollinating fig wasp foundress number (Fig. 3d). Taken together, these results suggest *Klebsiella* bacteria resembling *Klebsiella* sp. WOUb2 are commonplace in the natural environment of *C. inopinata*.

## Discussion

### A potentially ancient host/microbe interaction

Here, we screened a panel of microbes isolated from the natural environment of *C. inopinata* for their impacts on nematode population growth (Fig. 1). One isolate, *Klebsiella* sp. WOUb2, promotes fitness in *C. inopinata* (Fig. 1c) and *C. elegans* (Supplemental Figure 1) via variable life history mechanisms (Fig 2): it drives higher fecundity in *C. elegans fog-2(q71)* mutants (Fig. 2a, Supplementary figure 2a) but only faster development in *C. inopinata* (Fig. 2b, Supplementary figure 2b) at 25°C. And, sequence comparisons reveal species related to *Klebsiella* sp. WOUb2 are frequently found in the natural environment of *C. inopinata* (Fig. 3b-d). *Klebsiella* species are also found associated with fig wasps (deposited but unpublished sequences associated with *Ceratosolen solmsi* genome project (Xiao et al. 2013)), plant-associated insects (Adams et al. 2011; Vargas-Asensio et al. 2014), and *C. elegans* (Samuel et al. 2016; Bubrig et al. 2020). The observation that *Klebsiella* sp. WOUb2 drives these variable life history responses prompts the question—is this *Klebsiella*/*Caenorhabditis* relationship conserved across species? Multiple studies have shown *Klebsiella* bacteria are frequently encountered by *Caenorhabditis* nematodes, which is consistent with a conserved, ancient host/microbe relationship. *Raoultella* species (arguably synonymous with the genus *Klebsiella* (Y. Ma et al. 2021)) have previously been isolated from *C. elegans* (Dirksen et al. 2016; Samuel et al. 2016; Zhang et al. 2021) and *C. inopinata* (Kanzaki et al. 2018) in its natural environment, as well as from rotting plant substrates (Lee et al. 2020). And, as microbes of the *Enterobacteriaceae* family (of which *Klebsiella* is a member) are frequently found with *C. briggsae* and *C. remanei* (Dirksen et al. 2016; Johnke et al. 2025), *Caenorhabditis* nematodes have likely lived in close association with *Klebsiella*-like microbes since the emergence of the *Elegans* group over twelve million years ago (Fusca et al. 2025).

However, there is a possibility that this host/microbe relationship is not strictly conserved. While many studies have revealed *Klebsiella* microbes living in close association with *Caenorhabditis* nematodes, these same studies have found multiple wild *Caenorhabditis* samples where *Klebsiella* bacteria were not detected (Dirksen et al. 2016; Samuel et al. 2016; Johnke et al. 2025). That is, not *all Caenorhabditis* nematodes thrive with these microbes in their natural context. Moreover, in at least one study, *Raoultella* ASVs were more prevalent on substrates than in nematode microbiomes, potentially suggesting these bacteria are selected against within nematodes microbial communities (Supplemental Table S9 of (Johnke et al. 2025)). In addition, this and other studies (Vega and Gore 2017; Johnke et al. 2025), have suggested that individual nematode microbiome assembly is highly stochastic and driven by the presence of microbes on a given individual substrate. Thus, it is unclear just how frequently *Caenorhabditis* nematodes encounter and interact with *Raoultella*/*Klebsiella* bacteria in their natural context. Along these lines, *Klebsiella* bacteria were not detected in a modest sample of *C. briggsae* and *C. remanei* nematodes (Dirksen et al. 2016). Thus, it is also possible that the *Caenorhabditis*/*Klebsiella* associations observed among *C. elegans* and *C. inopinata* are coincidental, driven by stochastic processes, and do not represent a strictly conserved host/microbe relationship. Another scenario (that is not mutually exclusive) entails the recognition of functional redundancy among microbial species (Louca et al. 2018)— more closely related bacterial species are more likely to harbor similar sets of metabolic genes. It is then possible that while not all *Caenorhabditis* species encounter *Klebsiella* species in their natural context, they associate with a group of microbes that fill similar functional ecological roles. This is consistent with members of the *Enterobacteriaceae* family (of which *Klebsiella* is a member) being associated with the vast majority of *Caenorhabditis* microbiomes (including members of the “core” microbiome (Zhang et al. 2017)). Thus, even if this *Caenorhabditis*/*Klebsiella* relationship is not strictly conserved among species, it is reasonable to suspect that *Caenorhabditis* nematodes have been thriving in close association with functionally-similar microbes for millions of years.

### Klebsiella species modulate life history responses in nematodes

*Klebsiella* sp. WOUb2 promotes increased reproductive success in both *C. elegans* and *C. inopinata* (Fig. 2). Indeed, previous studies have shown that *Klebsiella/Raoultella* species influence reproductive success in these nematodes. *Raoultella* sp. CiN001 (with the most similar sequence to *Klebsiella* sp. WOUb2 detected in the NCBI nr database; Fig. 3A) increased population growth in *C. inopinata* (Kanzaki et al. 2018). *Raoultella* (formerly *Enterobacter*) sp. JUb54 increases the developmental rate of *C. elegans* (Samuel et al. 2016), similarly to our observations of wildtype *C. elegans* on *Klebsiella* sp. WOUb2 (Fig. 2b). Conversely, when *C. elegans* is grown on a complex bacterial community (called the BIGbiome), *Raoultella* sp. colonization of the *C. elegans* gut is correlated with *decreased* nematode developmental rate (Table S3 of (Zhang et al. 2021)). In addition, *C. elegans* prefers *Raoultella* sp. JUb38 to *E. coli* in behavioral preference assays (Sengupta et al. 2024). *Raoultella* sp. JUb54 also promotes increased dauer survival (Bubrig et al. 2020). And, as *C. elegans* is a biomedical model system, some studies have exposed *C. elegans* to the human pathogen, *Klebsiella pneumoniae*. Although likely not associated with its natural environment, *Klebsiella pneumoniae* has negative impacts on *C. elegans* such as reduced fecundity and lifespan (Kamaladevi and Balamurugan 2015; Yang et al. 2023). *Caenorhabditis* nematodes then certainly exhibit variable physiological, behavioral, and life history responses when reared on different *Klebsiella*/*Raoultella* isolates.

### fog-2 *and a Klebsiella-mediated increase in individual fecundity in* C. elegans

*fog-2* mutants had significantly higher individual fecundity when reared on *Klebsiella* sp. WOUb2 compared to *E. coli* irrespective of temperature (Fig. 2a, Supplementary figure 2a), and this response was not seen in wild-type *C. elegans* (Supplementary figure 3). One likely reason for this observation is that wild-type *C. elegans* fecundity measurements were not performed in the presence of males.

Wild-type hermaphrodites are sperm-limited (Ward and Carrel 1979; Hodgkin and Barnes 1991; Gimond et al. 2019), and microbe-mediated parental effects (both maternal and paternal) have been shown to modulate life history traits, behavior, and pathogen resistance in *C. elegans* progeny (Ha et al. 2010; Perez et al. 2017; Pereira et al. 2020; Willis et al. 2024; Pete and Hunter 2025). Thus, it is likely that wild-type *C. elegans* hermaphrodites reared on *Klebsiella* sp. WOUb2 would also demonstrate increased individual fecundity in the presence of males. Alternatively, differences in *fog-2* activity itself may be driving changes in microbe-dependent fecundity (independently of its male-promoting role). *fog-2* is a *C. elegans*-specific (Nayak et al. 2004), F-box encoding (Clifford et al. 2000) gene necessary for hermaphrodite-specific spermatogenesis (male spermatogenesis is unaffected (Schedl and Kimble 1988)). *fog-2* has also been implicated in sex pheromone production, although this is also thought to be intimately tied to its role in spermatogenesis (Leighton et al. 2014). F-box genes have undergone tremendous diversification among *Caenorhabditis* species, with high variation in interspecific copy number (Wang et al. 2021) and intraspecific nucleotide diversity (F. Ma et al. 2021). One explanation for this variation is that these genes may be implicated in innate immune responses (Singh and Luallen 2024). Although no known immunity role for *fog-2* has been established, *fog-2* may have some cryptic microbe-associated functions, which could explain the patterns we have observed here. If males do drive the patterns of *Klebsiella*-mediated increased fecundity in *fog-2* mutants (Fig. 2a, Supplementary figure 2-3), it is unclear why there is a lack of such a response in the obligately outcrossing *C. inopinata* (Fig. 2a). One possible explanation is that *C. elegans fog-2* mutants have increased fecundity because of some ancestral male-specific response to *Klebsiella* bacteria commonly found on microbe-rich rotting plant detritus (the ancestral environment of *Caenorhabditis* nematodes (Kiontke et al. 2011)). And, perhaps the shift to fresh figs, which are likely to be more microbe-limited than rotting plant detritus, has led to an evolutionary loss of a male-specific fecundity response to *Klebsiella* bacteria in *C. inopinata*. Alternatively, there may be some species-specific, cryptic feature of the *C. elegans* environment that is driving the evolution of this response only in this species. Future work specifically examining male phenotypic responses to these microbes across nematode species with variable reproductive modes will help to disentangle these possibilities.

### The evolution of divergent life history responses across beneficial interactions

Despite ultimately shared benefits in reproductive success on *Klebsiella* sp. WOUb2 (Fig. 1c; Supplemental Figure 1), *C. elegans* and *C. inopinata* nonetheless harbor variable life history responses to the same microbial strain (Fig. 2). If this host/microbe interaction is conserved (see above), then this suggests that such relationships can evolve along different evolutionary trajectories to promote such benefits via varying levels of investment into different life history mechanisms. That is, while the consequence remains the same, the proximate causal details are in flux. This phenomenon readily evokes a long-standing conjectural theory in host/pathogen co-evolutionary studies—the Red Queen hypothesis (Van Valen 1973; Clay and Kover 1996). Here, sustained antagonistic coevolution among pathogen and host at the molecular level underlies apparent stasis of phenotypes at higher levels of biological organization (Van Valen 1973; Clay and Kover 1996; Solé 2022). This idea has been used to explain myriad causes of host/pathogen evolutionary consequences such as those observed in *Daphnia* crustaceans and *Pasteuria* bacteria (Bourgeois et al. 2021); patterns of diversity in human immunity genes (Siddle and Quintana-Murci 2014; Ejsmond and Radwan 2015); and virulence profiles in rust fungi/poplar systems (Persoons et al. 2017).

However, the Red Queen hypothesis requires that divergent paths of *antagonistic* interactions are sustained to promote stasis, as novel alleles are constantly fixing in an evolutionary arms race between host defensive and pathogen offensive traits (Clay and Kover 1996; Paterson et al. 2010). Here, we have described a somewhat analogous case where hosts *benefit* from a microbial interaction via variable underlying mechanisms. While our observation evokes Red Queen dynamics because of these patterns of stasis and divergence across levels of biological organization, the valence of the microbial interaction on host fitness suggests that the evolutionary dynamics in play are likely to be quite different.

That is, Red Queen dynamics entail divergent selection, whereas this proposed scenario of a conserved beneficial interaction suggests stabilizing selection is at play. While molecular flux (reminiscent of Red Queen scenarios) is usually unexpected under stabilizing selection, such changes can be explained by compensatory evolution (Haag and Molla 2005; Haag 2007). Here, changes in the host that negatively impact the beneficial relationship can be rectified via compensatory changes in the microbe (or vice-versa). If different patterns of allelic substitution occur among divergent lineages, this could potentially lead to such patterns where fitness benefits are conserved while the underling life history causes have changed (Fig 1c; Supplemental Figures 1-3; Fig. 2a-b). Compensatory evolution has long been invoked in molecular evolution to explain sequence divergence despite functional conservation, and plausible examples of such change abound including the persistence of tRNA variation (Kern and Kondrashov 2004), compensatory changes to maintain 3D protein structure in hormone receptors (Ortlund et al. 2007), and allosteric changes that restore hemoglobin function (Natarajan et al. 2013). Moreover, a compensatory model has the potential to extend processes proposed to underlie such phenomena as Dobzhansky-Mueller incompatibilities (DMIs; (Kondrashov et al. 2002)) and developmental system drift (DSD; (True and Haag 2001)) to interactions among species. That is, both DSD and DMIs have been proposed to evolve via compensatory change under stabilizing selection to promote molecular divergence in the face of phenotypic stasis (True and Haag 2001; Kondrashov et al. 2002). Indeed, as *C. elegans* reproduces mainly by self-fertile hermaphroditism (Cutter et al. 2019), differences in host-microbe evolutionary outcomes are expected to diverge in comparison to an obligate outcrosser like *C. inopinata*, as sexual reproduction is thought to enable Red Queen-like dynamics (Morran et al. 2011). Here, conserved microbe-driven fitness benefits conferred via variable mechanisms in different lineages affords a potential analogous example that could extend such concepts to host/microbe interactions.

### The promise of a coevolutionary genetics of host-microbe interactions

We have described how a microbe isolated from the natural environment promotes reproductive success via variable life history mechanisms in two nematode sister species. *C. elegans* is a longstanding model system with a sophisticated experimental genetic toolkit (Nance and Frøkjær-Jensen 2019; Calarco et al. 2025) that has been use to address myriad questions across the wide scope of the biological sciences (Corsi et al. 2015). The discovery of its ecological niche (Kiontke et al. 2011), in tandem with the collection of hundreds of wild microbial (Dirksen et al. 2020) and *C. elegans* (Crombie et al. 2024) isolates, has opened the door to molecular ecological and quantitative genetics of host/microbe interactions. But beyond this, the exponential pace of *Caenorhabditis* species discovery (Sloat et al. 2022), the widespread applicability of CRISPR/Cas9 technology across nematode species (Du et al. 2021), and the increasing affordability of genome sequencing will likewise enable a molecular evolutionary genetics of host/microbe coevolution. These developments herald the promise of a research program interrogating the molecular substrates of such interactions. The work described above support such an optimistic horizon for future research in this area.

## Methods

### Isolation of fig microbes

To isolate bacteria from the natural environment of *C. inopinata, Ficus septica* figs were sampled in Taipei, Taiwan in 2019. Figs were dissected in 4 ml of sterile M9 buffer (Brenner 1974) in a sterile petri dish. 200 μl of fig suspension was transferred to a sterile LB plate and left at room temperature. Subsequently, monocultures were generated on LB media reared at room temperature. Stocks were preserved by adding 1.5 mL of a saturated liquid culture in LB to 0.5 ml of 60% glycerol; these were then frozen in liquid nitrogen and stored at −80°C.

### Preliminary identification of fig bacteria and genome assembly of Klebsiella sp. WOUb2

For DNA preparations, bacteria were grown in 2 mL LB liquid cultures. DNA was prepped with the DNeasy Powerlyzer PowerSoil Kit (cells were lysed with a Fisherbrand Bead Mill 4 for 10 minutes). Genus identities were determined via PCR and Sanger sequencing of the ITS region of the 16S rRNA gene (using the 27F and 1492R primers (Heuer et al. 1997)). The NCBI nr database was queried with these sequences with BLASTN (Camacho et al. 2009), and the genus of the top hit by bit score was used as for a preliminary taxonomic identification (list of all bacterial strains used in this study alongside their 16S sequences can be found in Supplemental Table Sheet 1). To better situate *Klebsiella* sp. WOUb2 in its phylogenetic context, the genome of *Klebsiella* sp. WOUb2 was assembled by Plasmidsaurus via Oxford Nanopore long-read sequencing (Supplemental Table Sheet 3). Phylogenetic methods and traditional microbial taxonomic approaches have revealed *Klebsiella* sp. WOUb2 to be a novel bacterial species (which will be described in a forthcoming manuscript; Miller et al. in prep).

### Nematode strains and maintenance

All nematodes were reared on peptone-free Nematode growth medium (NGM) plates. Peptone-free plates were used to control for the amount of available bacterial biomass on the NGM plates. All bacterial strains were reared in LB broth at 37°C to an optical density of 0.6 at 600nm. This, in conjunction with the peptone-free plates, was done to ensure that approximately the same amount of cellular material would be placed onto each experimental plate. For each experimental plate, 150ul of this bacterial solution was used for plate seeding. The *C. inopinata* strain NKZ35, *C. elegans* PD1074 (Brenner 1974), and *C. elegans* strain *fog-2(q71)* JK574 (Schedl and Kimble 1988) were used for all experiments in this work. Wild-type *C. elegans* strains are androdioecious (hermaphrodite/male), but *C. inopinata* is gonochoristic (female/male). Therefore, to account for differences in reproductive mode between these two species when measuring life history trait responses, the *C. elegans fog-2(q71)* JK574 mutant line was also used. This mutant line prevents spermatogenesis only in *C. elegans* hermaphrodites while it is maintained in males (Schedl and Kimble 1988), making the stocks functionally gonochoristic.

### Population growth

Synchronized *C. inopinata* NKZ35, *C. elegans* PD1074 and *C. elegans fog-2 (q71)* JK574 populations were generated through bleaching (Stiernagle 2006), and embryos acquired in this process were placed onto plates seeded with the respective experimental bacterial strain. These plates were incubated at 25°C until nematodes reached the L4 stage. For each cross of *C. inopinata* NKZ35 and *C. elegans fog-2 (q71)* JK574, four males and one female were placed onto a single plate seeded with a given bacterial strain. Single *C. elegans* PD1074 hermaphrodites were placed onto seeded bacterial plates for this assay. Nematodes were then allowed to proliferate for seven days, and the number of nematodes on each plate were counted. All estimates of population growth on a given fig bacterial strain were paired with controls where animals were reared on *E. coli* OP50. Failed crosses were those where either the L4 female died prior to reproduction or the number of nematodes at the end of one week did not exceed the initial amount (five nematodes: four males and one female). Crosses contaminated with non-specific bacteria were excluded.

### Individual fecundity and reproductive duration

Synchronized *C. inopinata* NKZ35, *C. elegans fog-2 (q71)* and *C. elegans* PD1074 embryos were generated through bleaching (Stiernagle 2006) and were raised on *Klebsiella* sp. WOUb2. or *E. coli* OP50 at 20°C or 25°C. Due to differences in developmental timing between *C. inopinata* and *C. elegans* (Kanzaki et al. 2018; Woodruff et al. 2019), the *C. elegans* populations were generated two days after the *C. inopinata* NKZ35 treatments. This allowed us to conduct the following experiment for all treatments at the same time (as the nematodes reached the L4 stage the same time despite their differences in developmental rate). When animals reached the L4 stage, crosses were performed with *C. inopinata* NKZ35 and *C. elegans fog-2 (q71)* JK574 to determine individual female fecundity (with one female and four males per plate). For *C. elegans* PD1074 L4 females, individual self-fertile nematodes were isolated for individual fecundity measurements. Every day, parents were transferred to a new plate, and the number of embryos and larvae generated on the previous plate were counted. For animals moved to a new plate, fresh males were also added (if needed for outcrosser strains) to maintain a consistent male:female ratio of 4:1. Parents were moved (and progeny counted) until no progeny were generated for three consecutive days (or the female died). The span from when reproduction began to when reproduction ended for a given individual female was recorded as the reproductive duration. In order to capture the total reproductive capabilities of each nematode reared on the different bacterial strains, animals that experienced matricidal hatching or died before the cessation of reproduction were excluded.

### Developmental timing and viability

Nematode populations (reared on either *Klebsiella* sp. WOUb2. or *E. coli* OP50) were synchronized by allowing gravid *C. inopinata* NKZ35 and *C. elegans fog-2 (q71)* JK574 females to lay embryos for six hours on the same respective bacterial strain. Females were removed, and plates were reared at 20°C or 25°C. The number of animals at each of four developmental stages (embryo, L1-L3 larva, L4 larva, and adult) were counted every day. Plates were counted until the number of adults did not increase for three consecutive days. The median time to reach every developmental milestone (L1-L3 larva, L4 stage, and adult stage) was inferred as in (Woodruff et al. 2019). Briefly, animals were coded as having reached or not reached a given milestone (for every day a given plate was measured). For each plate, a logistic model fit was generated (with the formula (milestone status ~ time)), and the time at which half of the nematodes were predicted to reach the milestone was noted (the median time to adulthood is reported in Fig. 2b). In addition to this, because the number of starting embryos and final number of adults was known for each plate, viability for each experimental treatment could be determined. Viability was measured as the fraction of embryos surviving to adulthood (Figure 2c).

### Body size

Using our developmental timing data (Figure 2b), we synchronized populations of *C. elegans fog-2 (q71)* and *C. inopinata* NKZ35 nematodes reared on *Klebsiella* sp. WOUb2. or *E. coli* OP50 so they would reach the young adult life stage at approximately the same time. Then, using 3% agarose in DI water as a mounting agent, we constructed microscope slides to take photographs of adult male and female nematodes from all experimental groups. Next, we pipetted 20 microliters of 0.35 mM Levamisole and placed it onto the 3% agarose mount. And, we placed nematodes from each sex-bacteria-nematode species treatment group onto distinct slides in the freshly pipetted Levamisole for each group. Using an Axioscope5 (Zeiss), we then took photographs of nematodes from each experimental group. Finally, using Fiji (Schindelin et al. 2012), we measured the lengths and widths of all nematodes in each experimental group after defining how many pixels corresponded to a given amount of microns.

## Acknowledgements

This work was supported by NSF award #2238788. Some strains were provided by the CGC, which is funded by NIH Office of Research Infrastructure Programs (P40 OD010440). Some of the computing for this project was performed at the OU Supercomputing Center for Education & Research (OSCER) at the University of Oklahoma (OU). We thank Jalynn Winrow for the contributing to the preliminary sequencing of some fig microbial isolates. We thank Paul Lawson and Samuel Miller for their work describing and phenotyping *Klebsiella* sp. WOUb2 and performing the phylogenetic analyses that determined this bacterial isolate is a novel species.

## Data availability

All data (raw/processed), supplementary material, and code can be accessed at the following url: (https://github.com/austincolelink/WOUb2_LifeHistory_October2025). The genome for *Klebsiella* sp. WOUb2 can be found on Genbank (SAMN41746641: WOUb02 (TaxID: 3161071)).

