## Supplemental Figures for "A novel, fig-associated microbe promotes reproductive success via variable life history mechanisms in *C. elegans* and *C. inopinata*"

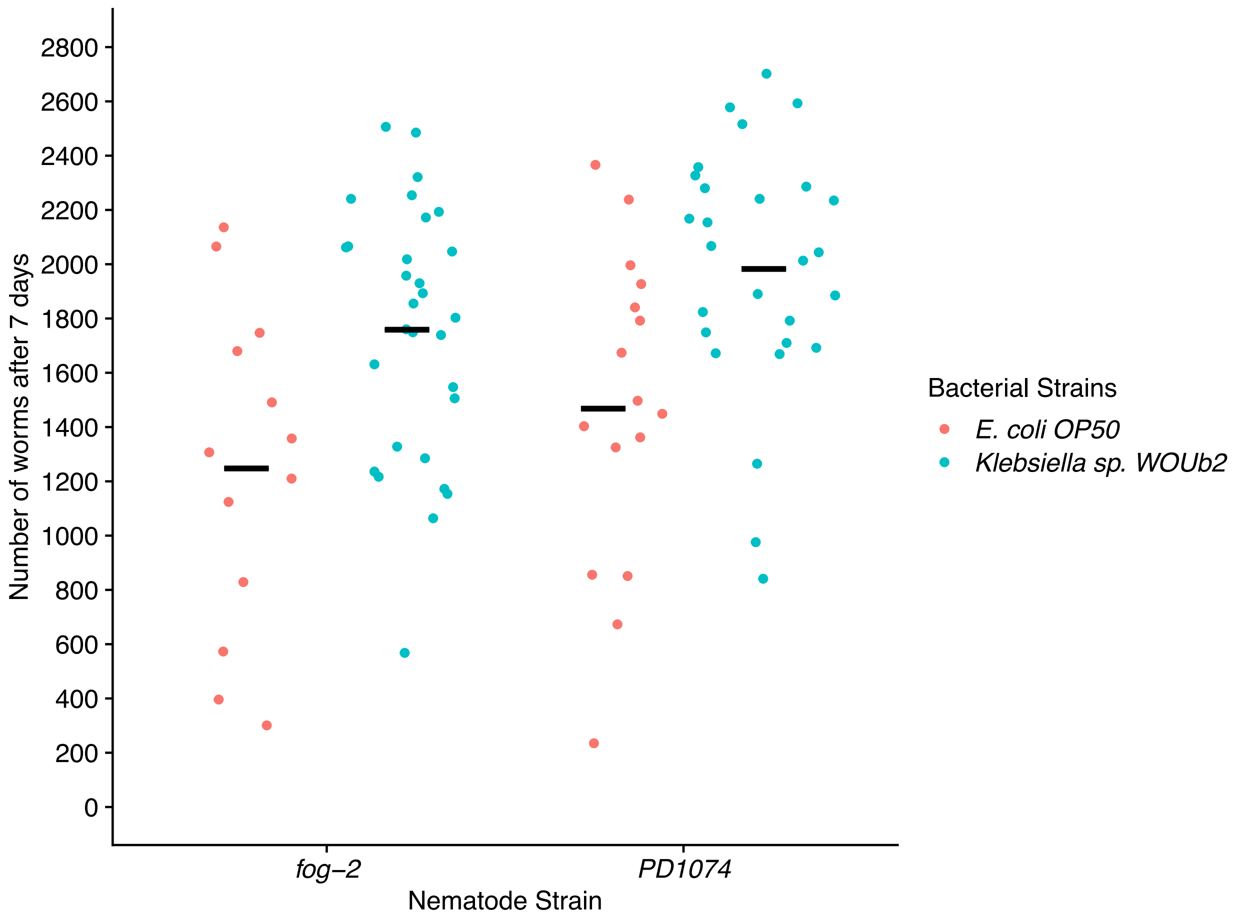


Supplementary Figure 1: *C. elegans* shows increased population growth reared on *Klebsiella sp.* WOUb2 compared to *E. coli* OP50. Each datapoint represents the number of nematodes on a single plate at the end of the seven-day period for a given bacterial strain. Crossbars represent means. Sina plots are strip plots that take the contours of a violin plot.

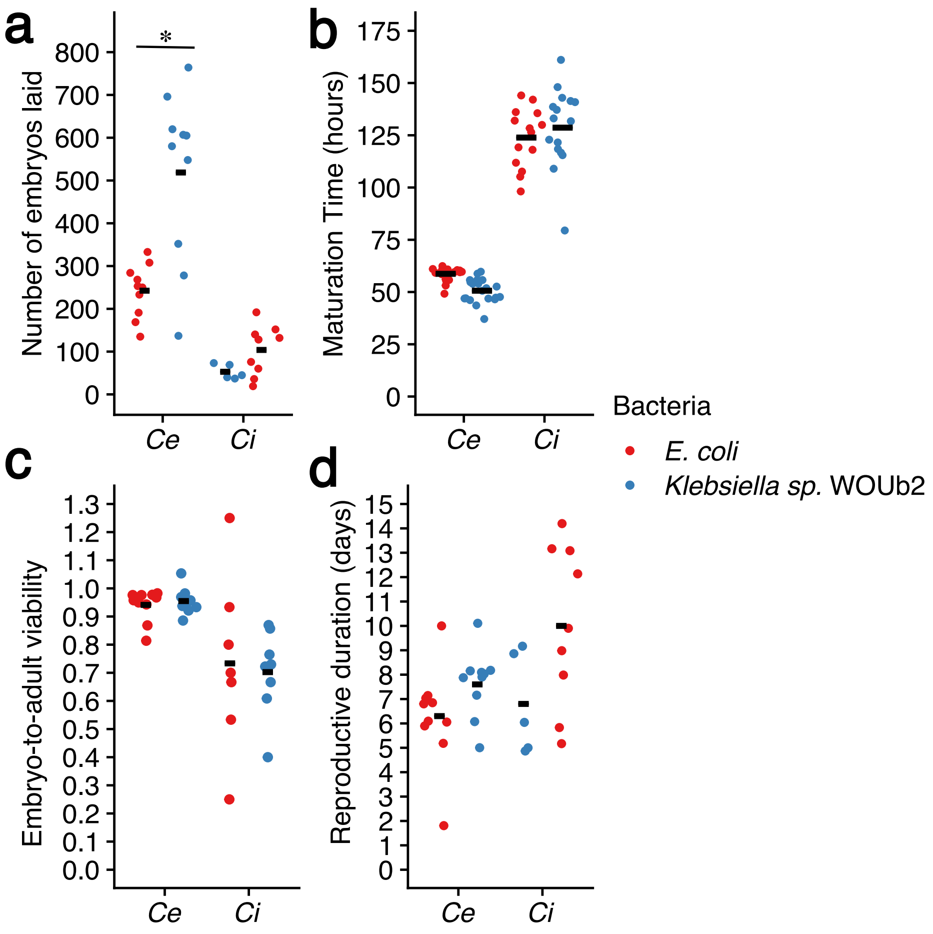


Supplementary Figure 2: Life history traits of *C. elegans* and *C. inopinata* reared on *Klebsiella sp*. WOUb2 and *E. coli* at 20C.

(a) Individual fecundity. *Klebsiella* sp. WOUb2 promotes higher individual fecundity in *C. elegans*, whereas in *C. inopinata*, individual fecundity is not changed. Each point represents the total number of embryos produced by an individual female.
(b) Developmental timing.
(c) Viability. Embryo-to-adult viability is unaffected for both *C. elegans* and *C. inopinata* in all rearing conditions. Each point represents the fraction of embryos surviving to adulthood.
(d) Reproductive period. *Klebsiella* sp. WOUb2 does not promote any major differences in reproductive duration among species. Each point represents the age of the last day of reproduction of an individual female.
For all plots, crossbars represent means. Sina plots are strip plots that take the contours of a violin plot.

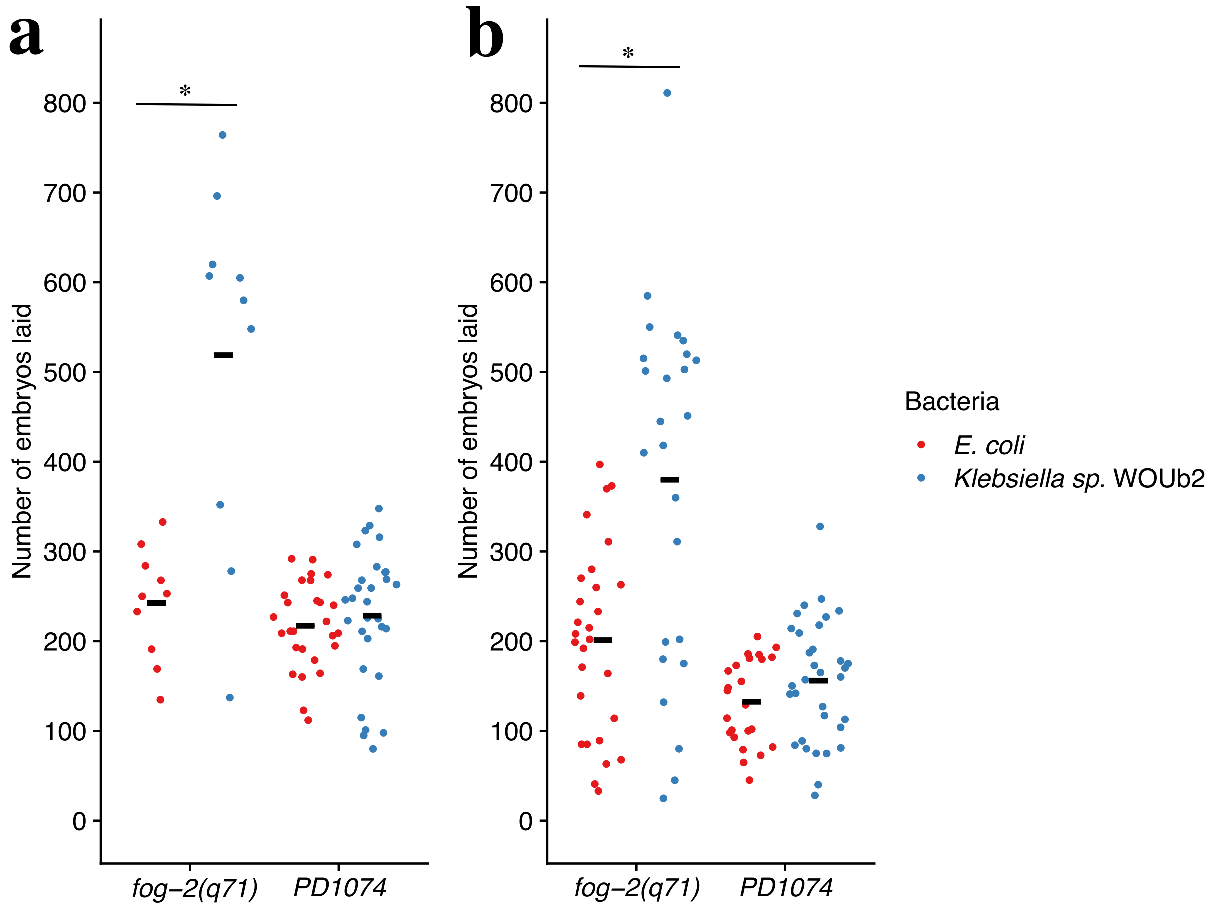


Supplementary Figure 3: Individual fecundity in *C. elegans* PD1074 WT vs. mutant *fog-2(q71)*

(a) 20C: *Klebsiella* sp. WOUb2 promotes higher individual fecundity in *fog-2(q71)*, whereas in PD1074, individual fecundity is not changed.
(b) 25C: *Klebsiella* sp. WOUb2 promotes higher individual fecundity in *fog-2(q71)*, whereas in PD1074, individual fecundity is not changed.

PD1074 is derived from the standard N2 wild-type isogenic line (Yoshimura et al. 2019).
